# Circular RNA identification using a genomic language model and a small number of authenticated examples

**DOI:** 10.64898/2026.03.04.709677

**Authors:** Kang Li, Weixu Wang, Jiaohao Jiang, Jing Deng, Jie Zhang, Shulan Qiu, Weixiong Zhang

**Author notes:** Equal contribution (Kang Li, Weixu Wang).

## Abstract

Genomic language models (gLMs) hold great promise for deciphering biological sequences, yet their effectiveness is hindered by the limited number of experimentally verified examples available for model training, a ubiquitous bottleneck for supervised machine learning. To overcome this challenge, we developed *circFormer*, the first gLM-driven approach for circular RNA (circRNA) identification. circFormer integrates curriculum learning with gLM fine-tuning: a Nucleotide Transformer model is first trained on a small set of validated circRNAs, the resulting model is used as a teacher to score ∼2.3 million noisy candidates, and the model is then fine-tuned with the noisy candidates along with their scores to improve prediction. Operating either as a standalone predictor or as a filter for existing pipelines, circFormer consistently outperformed traditional machine-learning approaches in accuracy and robustness. Among 50 circFormer-selected candidates that were overlooked by most existing tools, experimental validation using RNase R digestion and RT-qPCR confirmed 94.1% (32/34) of the evaluable candidates as genuine circRNAs. To enhance interpretability, we introduced a model-agnostic, dual-level explainable AI strategy that reveals mechanistic signatures of circRNA formation. circFormer provides a scalable, interpretable, and generalizable framework for converting noisy high-throughput data into reliable functional annotations, highlighting a practical path forward for gLM-based genomics in data-scarce settings.

## 1. INTRODUCTION

The application of deep learning to biology, particularly through large genomic language models (gLMs), has emerged as a powerful strategy for functional genomics[1-3]. This paradigm shift is driven by the rapid expansion of high-throughput sequencing, which produces large volumes of experimental data that enable the computational identification of millions of candidate functional elements, such as noncoding RNAs, regulatory regions, and splice sites[2, 3]. However, these large experimental datasets are often noisy, containing both genuine biological signals and artifacts introduced by experimental workflows or computational pipelines. In sharp contrast, experimentally verified ground-truth examples remain limited, posing a significant challenge for the effective application of gLMs that rely on abundant, high-quality labeled examples for model building, uncertainty calibration, and performance benchmarking.

The profound disparity between the scarcity of authenticated examples and the abundance of noisy, unlabeled data poses a major barrier to training and fine-tuning reliable gLMs. Models trained or fine-tuned on small, experimentally verified datasets are prone to overfitting and lack generalizability, whereas those trained on large, noisy, unlabeled datasets typically exhibit reduced predictive power and reliability. In both cases, model performance is constrained less by model complexity or data volume than by the limited availability of representative ground-truth examples.

The identification of circular RNAs (circRNAs) exemplifies this fundamental challenge that nearly all machine learning methods face. As covalently closed, single-stranded RNAs found across eukaryotes[4], circRNAs perform diverse regulatory functions, from interacting with the transcriptional machinery[5] to function as microRNA sponges[6, 7]. Although hundreds of thousands of putative circRNAs have been identified from RNA-seq data[8] and catalogued in databases[9], the majority of these candidates lack experimental validation[10]. The high cost and low throughput of gold-standard assays such as qPCR and Sanger sequencing severely constrain verification efforts, leaving the field dependent on two imperfect resources: small, high-quality datasets (often fewer than 1,000 verified examples) and vast, noisy datasets prone to false positives from repetitive genomic sequences, experimental artifacts, or genome mapping ambiguities. This ubiquitous data imbalance hampers gLM workflows that require large, reliably labeled datasets to adapt general genomic representations for circRNA identification, function analysis and performance benchmarking. Therefore, the scarcity of ground-truth examples remains the primary bottleneck to transforming gLMs into reliable tools for functional genomics.

To address the ubiquitous and fundamental challenge of insufficient authenticated data, we developed circFormer, a framework that integrates gLM methodologies with a curriculum-based learning strategy to effectively leverage unverified data. We demonstrate its effectiveness in identifying, validating and characterizing circRNAs. Using the Nucleotide Transformer[3] as a backbone pre-trained gLM, we first fine-tuned it on a small set of 939 experimentally validated circRNAs, and then refined the fine-tuned model with a second round of fine-tuning on 2.3 million noisy, unlabeled data. Furthermore, to fully leverage circFormer for circRNA investigation, we introduced a model-agnostic, dual-level explainable AI (xAI) approach to elucidate the model’s reasoning and gain mechanistic insights into circRNA biogenesis. Using *in silico* mutagenesis, we demonstrated that the model identified distinct sequence grammars associated with AG/GT and non-AG/GT back splicing. Crucially, to overcome the “black box” nature of deep learning, where internal representations are often dense and polysemantic, we adopted Sparse Autoencoders (SAEs)[11-13]. By decomposing the model’s entangled latent activations into a sparse, overcomplete set of interpretable features, SAEs enabled unsupervised isolation of specific biological signals directly from the high-dimensional embedding space. Together, the circFormer approach provides a generic, scalable framework for circRNA identification and interpretation, offering both methodological advances and deeper biological insight into RNA circularization mechanisms.

## 2. RESULTS

### 2.1. circFormer: A curriculum learning-empowered gLM to address the “data scarcity-noise” paradox

To address the challenge of the lack of authenticated examples, we developed a three-phase curriculum-learning approach (Fig. 1A) for fine-tuning a pre-trained gLM. Taking circRNA identification as a case study, we used the small 500-million-parameter Nucleotide Transformer (NT)[3], one of the best gLMs developed to date, as the pre-trained gLM. This small NT model was preferred because it slightly outperformed its larger counterparts (data not shown) and was adequate for circRNA discovery and analysis.

**Fig. 1.**
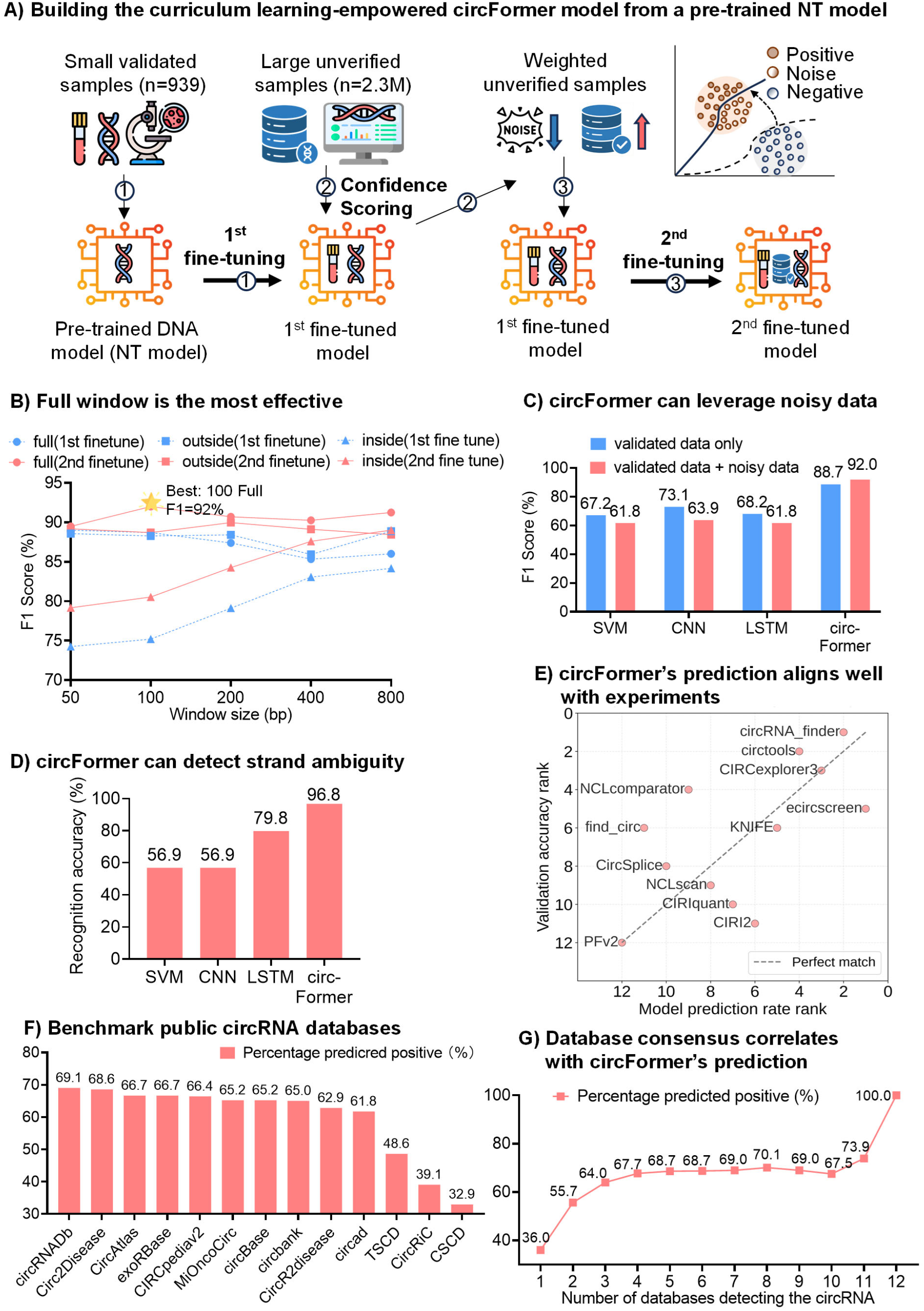
The circformer’s approach and performance on circRNA identification. **A)** Schematic view of the 3-step curriculum learning framework utilizing a genomic language model (the NT model) and a small number of authentic training examples, where the steps are numbered. The two fine-turning steps conceptually adjust the decision boundaries that separate genuine and artifactual back-splicing junctions of circRNAs (small figure in the upper right corner). **B)** circFormer’s performance across sequence lengths (or window sizes) and sequence contexts. Blue and red lines represent performance after the first and second fine-tunings, respectively, and triangles, squares, and circles denote inside context using junction-spanning sequences, outside context using flanking sequences, and both contexts, respectively. **C)** Performance comparison, measured by F1 scores, of circFormer and three machine-learning models, showing the effect of curriculum learning using large noisy, unlabeled data. **D)** Performance of four methods on noisy RNA-seq data with strand ambiguity. **E)** Concordance between circFormer’s ranking of 12 popular circRNA-finding methods (x-axis) and their ranking by the experimental benchmarking in [8] (y-axis). **F)** Benchmarking 13 circRNA databases measured by the percentage of entries predicted as genuine circRNAs by circFormer. **G)** Database consensus correlates with model prediction. The line plot shows the model’s positive prediction rate with respect to the number of independent databases that detect a circRNA candidate.

In Phase 1, we fine-tuned the chosen NT model on a small set of 939 experimentally verified circRNAs from the latest benchmarking study[8]. To facilitate performance comparison with other methods, we used cross-validation, splitting the gold-standard dataset of 939 circRNAs into three parts: 80% for fine-tuning (72% for training, 8% for validation) and 20% for testing. The fine-tuning yielded a refined model with an Area Under the Curve (AUC) of 0.891 and an F1-score of 0.887. Subsequently, in Phase 2, we employed this fine-tuned model to score 2.341 million unique, noisy, unlabeled candidates aggregated from 13 circRNA databases (Supplemental Table S1)[9, 14-25]. Because the fine-tuned model has presumably extracted pertinent circRNA features from experimentally validated circRNAs, it would score candidate back-splicing junctions that resemble genuine ones higher than those that do not. Finally, in Phase 3, we fine-tuned the model a second time using this massive, noisy dataset, weighting each unlabeled example’s contribution to the training loss by its model-derived confidence score (see Methods).

The sequences used for fine-tuning were genomic segments centered over or adjacent to putative circRNA back-splicing sites. We tested three types of sequences: the first from *k*-nt-long windows centered around the back-splicing sites (named full windows), the second from *k*-nt-long windows outside of but adjacent to circRNAs (named outside windows), and the third from *k*-nt-long windows inside of but adjacent to circRNAs (called inside windows). Note that the first and second window schemes cover the splicing signals for most circRNAs. We then compared these three schemes, varying *k* from 50 to 800 nt, at the end of Phase 1 and Phase 3 fine-tuning of the model (Fig. 1B). As expected, the third window scheme performed poorly because it did not use splicing signals. The best cross-validation performance (F1 = 0.920) was achieved with the full-window scheme and *k* = 100 nt (Fig. 1B). The second fine-tuning indeed improved performance, indicating that the model could learn genuine circRNA features from the noisy unlabeled dataset. This empirical analysis also suggested that both genomic segments inside and outside of circRNAs contribute to back splicing underlying circRNA formation.

Curriculum learning is an effective strategy for facilitating multiple rounds of fine-tuning, a unique feature of gLMs that conventional machine learning methods lack. To further assess this feature, we compared circFormer with standard machine-learning (SVM) and deep-learning (CNN and LSTM) methods (see Methods). For fair comparison, we trained two models for each of these three methods: one using only the small, authenticated dataset, and the other using both the small authenticated dataset and the large 2.3M noisy dataset. While all these models achieved comparable performance when trained on the small dataset, they failed to make any use of the large dataset; instead, their performance degraded due to data noise (Fig. 1C). In contrast, circFormer substantially outperformed all these conventional machine learning methods. More importantly, the curriculum-learning-enhanced circFormer improved its performance after the second-round fine-tuning using the large noisy dataset, increasing the AUC to 0.923 (a 3.2% improvement) and the F1-score to 0.920 (a 3.3% improvement; Fig. 1C; Supplemental Table S2).

Furthermore, we rigorously stress-tested the circFormer model against strand ambiguity, a pervasive source of false positives in RNA-seq data stemming from experimental errors or non-strand-specific RNA library preparation. We challenged the model with a “hard” negative set of “strand-contaminated decoys” (i.e., sequences derived from *bona fide* circRNAs but swapped to the opposite genomic strands). The circFormer model correctly rejected 96.8% of these decoys, improving on the Phase 1 baseline (96.3%) (Fig. 1D). This high discrimination accuracy confirmed that the model did not memorize genomic coordinates but internalized strand-specific sequence properties to robustly distinguish biological signals from technical artifacts.

### 2.2 circFormer distinguished false positives in public circRNA repositories

The circFormer model can run in a standalone mode to identify circRNAs from RNA-seq data processed by a genome mapper, e.g., the STAR method, which can nominate candidate back-splicing loci[26]. Alternatively, it can be used as a filter to remove false positives from circRNA-prediction methods. In the sequel, we substantiate this idea by assessing the quality of 12 of the 16 circRNA-detection methods that were experimentally evaluated in a recent large-scale benchmarking study[7, 8, 27-36] (Supplemental Table S3).

The remaining four methods lacking strand information were excluded, as no circRNA sequences could be determined. Specifically, we applied circFormer to evaluate the 12 pipelines and ranked them by the model’s calculated ‘authenticity rate’—the percentage of candidates predicted as true positives. This allowed testing whether circFormer could computationally replicate the experimental findings in [8].

Strikingly, circFormer’s rankings well correlated (Spearman’s ρ = 0.623, P = 0.03) with the experimental results in [8] that used qPCR, RNase R digestion, and amplicon sequencing (Fig. 1E). The existing methods with high performance by experimental validation, such as circRNA_finder (ranked 1), circtools (ranked 2), and CIRCexplorer3 (ranked 3), were similarly prioritized by circFormer. Conversely, tools that had lower validation rates, such as PFv2, were ranked lower by the model. This striking concordance demonstrates that circFormer effectively captures the intrinsic sequence features that distinguish experimental artifacts from genuine splicing events. It suggests that the model can serve as a reliable, automated computational benchmark for evaluating circRNA discovery software, potentially reducing the need for time-consuming and costly bench validation.

Encouraged by the high correlation between circFormer and experimental validation (Fig. 1E), we applied the model to score approximately 2.3 million unique putative circRNAs aggregated in 13 circRNA databases[9]. These databases were constructed using all or some of the 12 circRNA-finding methods recently benchmarked[8], each curating 306 to over 1.2 million human circRNAs (Supplemental Table S1). Out of the

2.3 million entries in the databases, circFormer retained 938,116 as true positives, filtering out more than half of the entries as likely to be transcriptional or splicing noise or technical artifacts. Consistent with this result, predicted validation rates were substantially heterogeneous across these repositories (Fig. 1F, Supplemental Table S1). Databases comprising manually curated or experimentally supported entries, such as circRNADb and Circ2Disease, achieved the highest validation scores of 69.1% and 68.6%, respectively (Fig. 1F). In contrast, comprehensive databases like CSCD and CircRiC showed significantly lower validation rates (Fig. 1F).

Furthermore, we assessed the agreement between circFormer’s predictions and the consensus of the existing methods. We ranked all circRNA candidates by the number of databases in which they appeared, showing a striking positive correlation between database consensus and the model’s prediction (Fig. 1G). Candidates unique to a single database—representing noises in most cases—were predicted as positive in only 36.0% of all cases. Conversely, as the number of supporting databases increased, the model’s validation rate rose to 100% for highly confident circRNAs across all 13 databases. This strong concordance indicates that circFormer successfully generalized to capture sequence-level features of stable, reproducible circRNAs, serving as a powerful *in silico* filter for high-throughput validation.

### 2.3. Experimental validation confirmed circFormer’s high performance

Beyond serving as an assessor of existing results and methods (Fig. E-G), circFormer also identified many high-confidence circRNAs that were consistently missed by most current pipelines. To confirm this observation, we experimentally validated circFormer’s top circRNAs in NCI-H23 lung cancer cells from [8] that were detected by no more than two existing methods. We focused on those that have not been empirically tested before and were longer than 60nt to facilitate PCR primer design. To be consistent with the latest large-scale experimental study in [8], we also considered circRNAs with both high and low expression abundance. Among the top 50 candidates we tested, 40 (80%) had at least 5 RNA-seq reads mapped to their back-splicing junctions (BSJ), and 10 (20%) had fewer than 5 BSJ-mapped reads (Supplemental Table S4). Note that these 50 candidates include 27 detected by exactly two of the 16 tools and 23 by only one, indicating that 87% (14 out of 16) of the existing methods have failed to detect them.

We subjected these 50 candidates to RNase R digestion followed by RT-qPCR, following closely the established benchmarking protocols in [8]. A candidate was deemed verified only if at least one of three untreated replicates exhibited a quantification cycle (*C*_*q*_) value of < 32 (see Methods). In the high-expression cohort, 28 of the 40 candidates met the evaluation criteria and all of them exhibited significant enrichment after RNase R treatment, confirming their circular structure. In the low-expression cohort, 6 were verified, of which 4 (66.7%) passed the RNase R validation (Supplemental Table S4). The reduced verification rate on the low-expression group is an expected outcome of the technical variance inherent to RT-qPCR, which disproportionately confounds the evaluation of low-abundance transcripts. In contrast, the 100% validation rate for high-expression group underscores circFormer’s exceptional precision when assessing transcripts within a reliable detection range. Overall, 32 of the 34 evaluable candidates (94.1%) were experimentally confirmed as genuine circRNAs. These results demonstrate that by internalizing sequence-level circularization signatures rather than relying on restrictive read-mapping heuristics, circFormer achieves superior sensitivity in detecting *bona fide* circRNAs that are systematically overlooked by conventional tools, suggesting that the gLM technology can substantially expand the annotated circRNA transcriptome.

### 2.4. CircFormer discovered intrinsic circRNA features

To maximize the utility of the circFormer model and augment it with model explainability, we introduced explainable AI (xAI) methodologies[37] to uncover the sequence features the model learned and used in its reasoning (Fig. 2A). The results increased the trustworthiness of the model and its results and shed new light on circRNA production mechanisms. We performed the analysis at both single-nucleotide and motif levels.

**Fig. 2.**
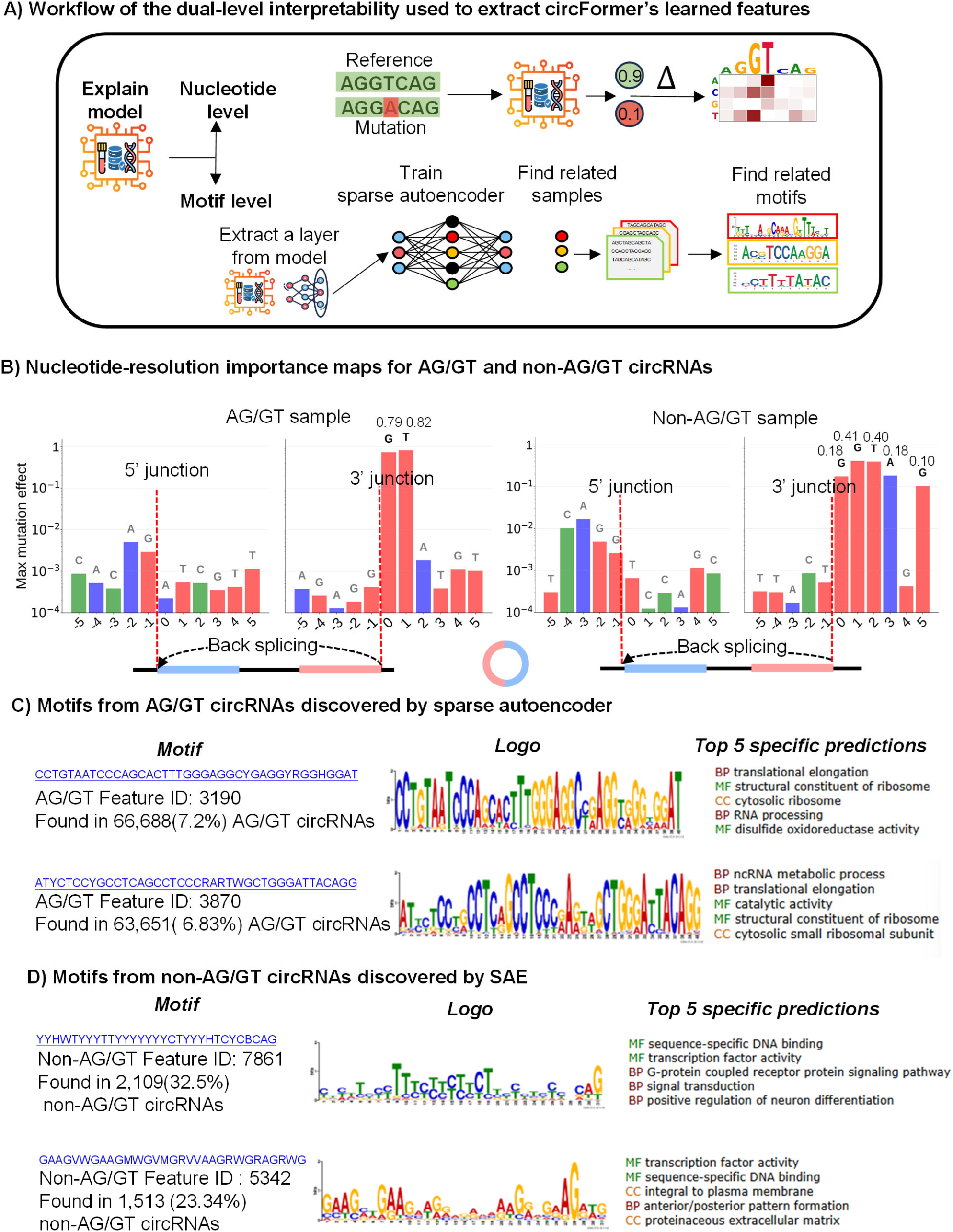
circFormer model interpretability and visualization of learned features. **A)** Schematic view of the dual-level explainable AI workflow. The diagram illustrates the two parallel approaches used to interrogate the features learned by the circFormer model: (1) Nucleotide level: *in silico* mutagenesis is applied to calculate the attribution score of each base by comparing the model’s output scores before and after mutation. (2) Motif level: Internal embeddings (hidden states) are extracted and processed by a Sparse Autoencoder (SAE) to isolate mono-semantic features. Highly activating sequences for these features are then analyzed using the MEME Suite to identify consensus motifs. **B)**. *In silico* mutagenesis profiles around back-splicing junctions. Bar charts show the maximum mutation effect (y-axis) at specific nucleotide positions relative to the 5’ and 3’ back-splicing junctions (x-axis) for representative AG/GT(left) and non-AG/GT (right) circRNAs. A corresponding schematic (bottom) depicts back-splicing junctions to indicate the genomic regions analyzed. The height of each bar represents the maximum divergence in prediction score induced by mutating the reference nucleotide to any of the other three bases. **C)** SAE-derived features for AG/GT circRNAs. Visualization of representative latent features (e.g., Features 3190 and 3870) extracted from the AG/GT dataset. Left: Consensus nucleotide sequence. Middle: MEME-generated sequence logo representing the position-specific probability matrix. Right: Top five Gene Ontology (GO) terms associated with the feature. **D)** SAE-derived features for non-AG/GT circRNAs. Visualization of representative latent features (e.g., Features 7861 and 5342) extracted from the non-AG/GT dataset, with a layout identical to (C).

#### 2.4.1. In silico mutagenesis at nucleotide resolution revealed the model’s focus on splice signals

We first employed *in silico* mutagenesis (ISM)[38] to quantify the importance of individual nucleotides for the model’s predictions (Fig. 2A). Briefly, ISM intentionally mutated a single nucleotide and tested whether and how the model’s prediction changed (see Methods). Because back splicing is a known mechanism of circRNA production, we grouped the validated 939 circRNAs into samples with AG/GT splice signals (90.3% of the total) and those without such signals. For AG/GT circRNAs (Fig. 2B, two left panels), ISM revealed two distinct patterns. In many samples, the model’s predictions were highly sensitive to mutations at the ‘GT’ donor site, with single-base changes reducing the probability of a true positive to below 0.8. The ‘AG’ acceptor site was also important, but often to a lesser degree, which may reflect the biological tolerance for minor splice signals (e.g., ‘AT/GT’). In other samples, mutations at the ‘AG-GT’ sites had little impact (score change < 0.003), suggesting the model had learned alternative, compensatory features elsewhere in the sequence to confirm the circRNA’s authenticity (Supplemental Fig. S1A). Furthermore, we examined the impact of consecutive mutations at one to four sites on model predictions. The regions with the greatest impact from mutations were concentrated near the ‘GT’ (Supplemental Fig. S1B).

For non-AG/GT circRNAs (Fig. 2B, two right panels), the model learned distinct sequence features. In some samples, the model’s predictions were highly sensitive to single-nucleotide changes. However, instead of focusing on the two-base-pair ‘GT’ site, the model’s attention was distributed over a wider region, often over a 4-base-pair (or larger) motif at the 3’ junction (Fig. 2B, two right panels). This provided evidence that the model did not simply memorize the ‘AG-GT’ rule but successfully learned distinct, alternative sequence patterns that define back-splicing.

#### 2.4.2. Sparse autoencoder uncovered mono-semantic features corresponding to known splicing motifs

To bridge the gap between the importance of single nucleotides and higher-order biological mechanisms, we employed a mechanistic interpretability approach. While the ISM method provided a high-resolution map at single-nucleotide resolution—highlighting discrete positions critical for prediction (e.g., splice sites; Fig. 2B—it failed to capture the cooperative logic of longer sequence patterns. To address this issue, we trained a sparse autoencoder (SAE) on circFormer model’s internal representations (Fig. 2A). The SAE was designed to decompose the dense, polysemantic 768-dimensional embeddings of the NT model into a high-dimensional dictionary of 12,800 “mono-semantic” features, where each latent feature ideally corresponds to a single, interpretable biological concept.

We first validated the SAE’s capability to disentangle these complex genomic representations. The SAE achieved rapid convergence during training and maintained high reconstruction fidelity, with R^2^ scores near 0.8 (Fig. S1C).

To systematically delineate the distinct regulatory logics underlying conventional (AG/GT) and atypical (non-AG/GT) circRNA biogenesis, we performed a comparative analysis to identify features preferentially activated in each specific class (see Methods)[39]. This approach revealed a striking dichotomy in the sequence determinants governing these two classes (Fig. 2C).

In AG/GT circRNAs, the regulatory landscape is dominated by motifs consistent with standard splicing machinery and translational regulation. Specifically, Feature 3190 (consensus: 5’-CCTGTAATCCCAGCACTTTGGGAGGCYGAGGYRGGHGGAT-3’), identified as an AG/GT-specific marker, was present in 7.2% of AG/GT sequences (n=66,688) (Fig. 2C, left panel; Fig. S1D Top panel). Similarly, Feature 3870 (consensus: 5’-ATYCTCCYGCCTCAGCCTCCCRARTWGCTGGGATTACAGG-3’) showed high prevalence (6.8%, n=63,651) (Fig. 2C, left panel; Fig. S1D Top panel). Functional annotation of these motifs via GOMo (Gene Ontology for Motifs)[40] revealed significant associations with translational elongation, cytosolic ribosome constituents, and RNA processing (Fig. 2C, right panel), suggesting that AG/GT circRNA biogenesis is coupled with cotranscriptional processing and translational machinery, supporting the notion that some circRNAs can be translated into proteins[41, 42].

Conversely, non-AG/GT circRNAs exhibited a fundamentally distinct motif composition, characterized by high-frequency motifs that were largely absent in the conventional AG/GT circRNAs. The most prominent motif, Feature 7861, comprising a pyrimidine-rich tract (consensus: 5’-YYHWTYYYTTYYYYYYYCTYYYHTCYCBCAG-3’), was detected in 32.5% of non-AG/GT sequences (n=2,109) yet appeared in only 5.3% of AG/GT sequences, indicating a significant enrichment in the non-AG/GT set (Fig. 2D, left panel; Fig. S1D, bottom panel). Furthermore, Feature 5342, a purine-rich motif (consensus: 5’-GAAGVWGAAGMWGVMGRVVAAGRWGRAGRWG-3’), was exclusive to the non-AG/GT class with 23.3% (n=1,513) prevalence (Fig. 2D, left panel; Fig. S1D, bottom panel). Crucially, the functional ontology of these atypical motifs diverged sharply from the RNA-processing themes observed in AG/GT sequences. The non-AG/GT-specific signatures were enriched for terms related to sequence-specific DNA binding, transcription factor activity, and integral plasma membrane components (Fig. 2D, right panel). This unexpected association suggested that non-AG/GT circRNAs might originate through alternative biogenesis pathways distinct from the classical spliceosome-mediated back-splicing, potentially involving specific transcription factor recruitment or coupling with membrane-associated signaling processes.

## 3. DISCUSSION

We have presented a novel approach that addresses a fundamental and challenging bottleneck in computational biology: the scarcity of gold-standard training data amidst an abundance of large, noisy datasets. This data dichotomy—where high-confidence experimentally validated examples are severely limited while computationally derived data are vast but artifact-prone—has long hindered the applications of machine-learning approaches, including foundation models, to biological and medical problems. Training exclusively in small experimentally validated datasets risks overfitting and poor generalization, whereas relying on large, noisy datasets propagates experimental and computational errors and false positives. Our curriculum learning-empowered genomic language model (gLM) approach provided a generalizable framework for overcoming this critical limitation by leveraging small, high-confidence datasets to guide robust learning using massive, unreliable data sources.

We demonstrated the utility and rigor of this novel framework in a case study that identified and validated circRNAs from large volumes of noisy RNA-seq data using a small number of high-quality training examples. circFormer is the first gLM-based approach for circRNA identification, achieving superior performance and outperforming existing methods. circFormer’s excellent discriminative power in distinguishing genuine from spurious back-splicing junctions can be attributed to two important factors. The first is the NT model, which was pre-trained on a large amount of genomic data from multiple species. The second, and more important, factor is the curriculum-based learning strategy that distills genuine biological signals – i.e., authentic back-splicing junctions – from an enormous amount of noise, including sequencing errors, genome-mapping artifacts, and irrelevant biological signals such as linear splicing. This learning strategy first leveraged a small set of 939 real circRNAs to fine-tune the NT model to extract intrinsic features of back splicing. It then used the refined model as a teacher to score the noisy samples. Finally, it used the loss function scores from the second round of fine-tuning to define a new discriminative boundary that more effectively separates authentic from spurious back-splicing junctions. The gLM methodology and curriculum learning strategy are thus perfectly integrated to overcome the fundamental challenge of insufficient authentic training examples that nearly all machine-learning methods face when applied to genomics, biology and medicine.

A key contribution of this study is exposing the inherent biases in current heuristic-based discovery pipelines, which typically rely on rigid alignment rules and consensus filtering. While these methods converge on a high consensus but limited set of annotations, they collectively neglect a vast unannotated repertoire of circular RNAs that deviate from predefined patterns. In contrast, circFormer’s predictive power transcends these computational constraints, as evidenced by our rigorous orthogonal wet-lab validation. Using RNase R digestion and RT-qPCR, we confirmed that 32 of 34 tested candidates (94.1%) were *bona fide* circular transcripts, despite being consistently missed by most 16 mainstream discovery tools. Notably, circFormer achieved a 100% validation rate (28/28) among candidates in the high-expression cohort, demonstrating exceptional precision when transcript abundance allows for reliable qPCR evaluation. The lower validation rate (66.7%, 4/6) in the low-expression cohort reflects established technical limitations of qPCR in assessing low-abundance transcripts, rather than a deficit in model prediction. These authentic yet “low-consensus” molecules represent a hidden layer of transcriptomic complexity captured by internalizing intrinsic sequence-level signatures rather than relying on read-mapping heuristics. To ensure practical utility, we developed circFormer-STAR, a streamlined pipeline that integrates our foundation model with the industry-standard STAR aligner to reduce false-discovery rates.

Beyond establishing a robust predictive tool, our study addressed a critical challenge in genomic AI: transforming high-performing “black boxes” into transparent engines for mechanistic understanding. By introducing a dual-level explainable AI (xAI) approach, we demonstrated that our circFormer model can transition from conventional statistical pattern matching to capturing the fundamental biological logic of RNA circularization. Importantly, the model autonomously deconvoluted the sequence determinants of conventional versus atypical RNA circulation without prior biological knowledge. For conventional (AG/GT) circRNAs, the model rediscovered the canonical splicing rules, associating them with motifs linked to ribosomal machinery and translational elongation. This serves as a rigorous “proof of concept” that the model has internalized the established rules of nuclear pre-mRNA processing. More distinctively, the model generated novel hypotheses regarding atypical (non-AG/GT) circRNAs. Unlike the splicing-centric logic of conventional RNA circulation, the model revealed that atypical circRNAs are governed by a distinct regulatory lexicon—rich in purine/pyrimidine disparities—associated with sequence-specific DNA binding, transcription factor activity, and integral membrane components. This divergence suggests that atypical circRNA biogenesis may not be merely a “splicing error” but a regulated process coupled with specific transcriptional programs or membrane-associated signaling pathways. Thus, a properly trained gLM can serve as an *in-silico* biologist, capable of both validating known paradigms and generating testable hypotheses for unexplored genomic features.

The circFormer approach can be improved. First, the model was fine-tuned on human data, limiting its immediate generalizability to other species. When applied to other species, it can be fine-tuned again using species-specific examples. Note that the curriculum learning strategy is species-independent and generalizable to other organisms. Second, the 100-nt window, while optimal for predictive performance, inherently limits the model’s ability to discover long-range regulatory elements distal to back-splicing junctions. Future work could explore model architectures that process longer sequences to capture these distal interactions.

## 4. MATERIALS AND METHODS

### 4.1. Data used

We constructed two distinct positive datasets: an experimentally validated, “gold-standard” set and a large but “noisy” set. The gold-standard dataset comprised 939 circRNAs experimentally validated by RNase R digestion, qPCR, and amplicon sequencing in the recent benchmarking study.[8] The noisy dataset contained 2,341,000 mixed real and putative circRNAs compiled from 13 public circRNA databases (Supplemental Table S1), most of which were computationally detected from RNA-seq data. All back-splicing junction sequences were extracted uniformly from the hg38 reference genome.

For each of two positive datasets, we constructed a negative sample set with a 1:1 positive-to-negative ratio. Each negative set consisted of two types of artifacts: (i) 20% “strand-flipped” artifacts, i.e., the reverse complementary sequences, of the circRNAs in the corresponding positive set, and (ii) 80% “random-junction” artifacts, generated by randomly selecting two exon/intron boundaries from the same genes to serve as 5’ and 3’ back-splicing points. All generated negative samples were filtered against the existing databases to exclude potential true positives.

### 4.2. Fine-tuning the NT model using curriculum learning

Taking the Nucleotide Transformer (NT) model[3] with 500M parameters as the pre-trained model, we fine-tuned it using a three-phase curriculum learning to develop circFormer.

#### Phase 1 – Fine-tuning using the gold-standard dataset

The 939 authentic circRNAs from [8] (positive sample) and an equal number of negative samples were split into training (72%), validation (8%), and test (20%) subsets. The pre-trained NT model was fine-tuned on the training set using Low-Rank Adaptation (LoRA) (r=1, lora_alpha=32, target_modules=[“query”, “value”]) for 5,000 steps, with a learning rate of 5e-4 and a batch size of 32 (adjusted for sequence length). The model checkpoint with the highest AUC on the validation set was saved. During training, binary cross-entropy loss was computed between the model’s logits and ground-truth labels. The predicted class label was determined by: label = 1 if P(positive) > 0.5, otherwise label = 0 (P(positive)+P(negative)=1).

#### Phase 2 – Confidence prediction for scoring

The fine-tuned model from Phase 1 was used as a teacher to score all 2.341 million putative circRNA samples in the noisy dataset, generating a confidence score – the probability of being a true positive – for each sample. Specifically, for each sample, the model generated class probabilities via softmax activation over the output logits. The confidence score was defined as the probability of the positive class (confidence = P(positive)).

#### Phase 3 – Curriculum-learning-based fine-tuning

The model from Phase 1 was further fine-tuned using both the gold-standard set (confidence score 1.0) and the noisy dataset (Phase 2 confidence scores). Samples from the noisy dataset were weighed in the cross-entropy loss function based on their predicted confidence scores from Phase 2, using a 5-tier system (confidence >=0.95: weight=1.0; 0.90 <= confidence <0.95: weight=0.8; 0.80 <= confidence <0.90: weight=0.6; 0.70 <= confidence <0.80: weight=0.4; confidence <0.70: weight=0.2). This discretization provides a simple, interpretable framework for progressively upweighting high-confidence samples while avoiding complete exclusion of samples (a minimum weight of 0.2 retains long-tail information and reduces selection bias). The tier thresholds (0.95/0.90/0.80/0.70) represent coarse-grained partitions of the confidence spectrum, ranging from “highly certain” to “potentially mislabeled,” with linear weight decay (1.0 to 0.2) balancing noise robustness against data utilization. This model was trained for 6,000 steps with a learning rate of 2e-4 and a gradient accumulation of 4. The final model was selected based on the best AUC on the gold-standard validation set.

All training was conducted using the Hugging Face transformers and peft libraries in PyTorch.

### 4.3. Model evaluation and baseline implementation

#### 4.3.1. Window-size evaluation

Using the gold-standard dataset, we tested five window sizes [50, 100, 200, 400, 800 nt] and three sequence contexts [‘full’ (the flanking and inside of a back-splicing junction), ‘inside’ (junction-adjacent), ‘outside’ (flanking only)]. Model performance for each configuration was assessed using 3 repeats of 5-fold stratified cross-validation.

#### 4.3.2. Baseline methods for comparison

To benchmark the performance of the circFormer approach, we implemented three conventional machine-learning and deep-learning methods: a Support Vector Machine (SVM), a Convolutional Neural Network (CNN), and a Long Short-Term Memory (LSTM) network. All models were trained and evaluated using a repeated stratified cross-validation scheme (5 folds, 3 repeats) to ensure robust performance estimation.

##### Support Vector Machine (SVM)

Input sequences were vectorized using k-mer frequency encoding with k=6. The resulting feature vectors were standardized using Z-score normalization. We employed a Linear SVC (Support Vector Classification) with a maximum of 10,000 iterations for convergence.

##### Deep Learning methods (CNN and LSTM)

For the deep learning baselines, nucleotide sequences were one-hot encoded into 4*xL* matrices, representing the four nucleotide channels (A, C, G, T) and a window of *L*=200-nt sequence around a back-splicing junction. Ambiguous bases (‘N’) were zero-padded. Both models were implemented in PyTorch and trained with the Adam optimizer (learning rate = 0.001) and Cross-Entropy loss for 10 epochs, with a batch size of 32.

- ***CNN Architecture***:The network comprised two 1D convolutional layers: the first with 64 filters (kernel size=7, padding=3) and the second with 128 filters (kernel size=5, padding=2). Each convolution was followed by batch normalization and ReLU activation. A global adaptive max-pooling layer was applied after the second block to extract salient features for the final fully connected classification layer.
- ***LSTM Architecture***: The LSTM model employed a bidirectional architecture to capture sequential dependencies from both forward and backward directions. It consisted of a single bidirectional LSTM layer with a hidden dimension of 128. The output from the final step was concatenated and fed into a fully connected layer (input dimension 256) to generate the prediction logits.

### 4.4. Nucleotide-level interpretation by *in silico* mutagenesis

*In silico* mutagenesis (ISM) was used to quantify the contribution of individual nucleotides to model predictions. For a given sequence of length *L* (e.g., 200 nt for the 100 nt full-window model), we generated 3 *x L* mutated variants by systematically substituting the nucleotide at each position with the other three nucleotides. The effect of each mutation was quantified as the difference between the original model’s positive-prediction score and the mutated sequence’s score. This procedure yielded an *L* × 4 attribution matrix featuring the impact of every possible single-nucleotide variant (SNV) on the model’s output.

### 4.5. Motif-level interpretation by sparse autoencoder

To systematically dissect the learned sequence features encoded in the circFormer model, we employed a Sparse Autoencoder (SAE) approach with Top-K sparsity constraints.[12] This approach addresses the fundamental “polysemanticity” problem in deep neural networks, in which individual neurons frequently encode multiple, functionally disparate biological motifs due to representational capacity limitations imposed by fixed-dimensional constraints.

#### 4.5.1. Feature extraction and training data preparation

To construct the training dataset for the sparse autoencoder, we compiled a corpus comprising the 939 experimentally validated samples and high-confidence model-predicted circRNAs from the 13 databases. To ensure balanced representation of AG/GT and non-AG/GT splice site architectures, we implemented a stratified sampling strategy: all non-AG/GT circRNA candidates (n=6,482) were retained, and an equal number of AG/GT examples were selected, ranked by prediction confidence. Splicing site classification incorporated a ±1 nucleotide positional tolerance window and recognized both AG/GT and AC/CT (reverse complementary of AG/GT) dinucleotide pairs as known major splicing signals, consistent with established splicing biology. For the resulting 12,964 sequences, we extracted the corresponding 768-dimensional hidden-state representations from Layer 11 of the fine-tuned model.

#### 4.5.2. Sparse autoencoder architecture and optimization

The SAE was engineered to decompose the dense, polysemantic input *x* ∈ ℝ^768^ into an overcomplete, highly dimensional dictionary of latent features *f* ∈ ℝ^12800^, representing an expansion factor of approximately 16×. This expansion facilitates the disentanglement of superimposed signals into “mono-semantic” features, where each latent direction corresponds to a distinct, interpretable biological motif. The architecture consists of:

- ***Encoder***: A linear transformation *W*_*enc*_ followed by Top-K activation to map dense inputs : *x* ∈ ℝ^*768*^ to sparse latent coded: *f* ∈ ℝ^*12800*^, *f* = *TopK*(*W*_*enc*_ ∗ *x* + *b*_*enc*_, *k* = 64)where only the 64 most strongly activated features were retained per example, with all others set to zero.
- ***Decoder***: A linear transformation was used to reconstruct the original input 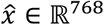 from the sparse code: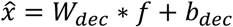
- ***Sparsity Constraint***: Unlike standard autoencoders, we enforced strict sparsity, ensuring that only a small fraction of features was active for each input. The training objective minimized a composite loss function:

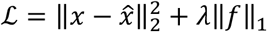

where the first term ensures reconstruction fidelity (preserving biological information) and the second term (L1 penalty, *λ =* 0.001) enforces sparsity. The model was trained with the Adam optimizer, with performance evaluated using reconstruction scores.

#### 4.5.3. Feature discovery and biological interpretation

To comprehensively characterize the learned feature dictionary, we conducted a three-pronged analysis identifying features with distinct activation patterns:

- ***Globally Dominant Features*** (n=10): These sequences represent archetypal inputs for specific features. Features exhibiting maximal mean activation across all sequences, representing ubiquitous structural or compositional motifs.
- ***AG/GT-Specific features*** (n=10): Features preferentially activated by circRNAs with AG/GT, encoding prototypical splicing consensus sequences. (Δ*activation = mean*_*AG*/*GT*_ *-mean*_*non-AG*/*GT*_)
- ***Non-AG/GT-Specific Features*** (n=10): Features demonstrating significantly elevated activation in non-AG/GT ones relative to AG/GT ones (Δ*activation = mean*_*non-AG*/*GT*_ *-mean*_*AG*/*GT*_), capturing sequence signatures unique to atypical splicing architectures.

This strategy yielded 23 unique features after de-duplication across categories for downstream mechanistic interrogation.

#### 4.5.4. Motif discovery by MEME suite

For each selected feature, we performed systematic motif extraction to decipher the sequence determinants driving its activation:

- ***Maximal Activation Sampling***: For each feature f_i, we identified the top 50 sequences eliciting the highest activation values, representing archetypal examples of that feature’s encoded motif.
- ***Motif Discovery***: These top 50 sequences were subjected to *de novo* motif analysis using MEME[43] (v5.5.9) with the following parameters: DNA alphabet, zero-or-one occurrence per sequence model (ZOOPS), classic objective function, motif width range 5-40 nucleotides, and first-order Markov background model.
- ***Validation***: Discovered motifs were summarized as Position Weight Matrices (PWMs) and sequence logos, and statistical significance was assessed using E-values. Features yielding significant motifs (E-value < 0.05) were retained for biological interpretation.

This integrated computational pipeline enables systematic mapping between abstract feature representations in the model’s latent space and interpretable, biologically grounded sequence motifs, thereby elucidating the mechanistic basis of the model’s classification decisions.

### 4.6. circFormer-STAR: A gLM-enhanced circRNA identification pipeline

To facilitate the adoption of our approach in standard bioinformatics workflows, we developed circFormer-STAR, an integrated command-line tool designed for high-precision circRNA discovery. Unlike conventional pipelines that rely solely on heuristic filters, our tool directly interfaces with the STAR aligner—a gold standard in RNA-seq processing.

Specifically, circFormer-STAR takes the raw Chimeric.out.junction file produced by STAR as input, extracts the genomic sequences flanking the candidate back-splicing junctions and applies the circFormer model to score each candidate’s authenticity. This integration bridges the gap between raw sequence alignment and sophisticated deep-learning discrimination, providing a streamlined “alignment-to-annotation” solution. To support open science and reproducibility, the entire circFormer framework, including the pre-trained models and the STAR integration scripts, is open-sourced and available at https://github.com/GenomicMedicine/circFormer.

### 4.7. Experimental verification: RT-qPCR and RNase R treatment

RNase R digestion was performed followed by reverse transcription quantitative PCR (RT-qPCR). Total RNA was extracted and treated with RNase R (Adamas-life, Shanghai, China) to degrade linear RNA species. Matched untreated controls were processed in parallel under identical conditions, except for omitting the enzyme. Following digestion, the RNA was purified and reverse-transcribed into cDNA using a reverse transcription kit (Vazyme, Nanjing, China) according to the manufacturer’s protocol.

RT-qPCR was performed using Vazyme SYBR Green-based reagents on a real-time PCR detection system. Each circRNA candidate was assayed in technical triplicate for both RNase R-treated and untreated conditions. To ensure a rigorous and standardized assessment, all validation criteria were adopted directly from the comprehensive benchmarking study.[8] A candidate was deemed evaluable only if at least one of the three untreated replicates yielded a quantification cycle (*C*_*q*_) value of < 32. Candidates failing to meet this threshold across all untreated replicates were excluded from downstream analysis, as their low baseline expression precludes an accurate assessment of RNase R resistance. For all tested candidates, we applied a conservative “best-case” strategy to calculate the cycle shift Δ(*C*_*q*_) to rigorously account for technical variability. Specifically, (*C*_*q*_) was calculated as the difference between the minimum (*C*_*q*_) of the treated replicates and the maximum (*C*_*q*_) of the untreated replicates:

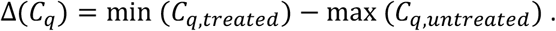

Based on the criteria, circRNAs exhibiting a Δ(*C*_*q*_) ≤ 3 cycles were considered resistant to RNase R digestion and considered verified true circRNAs. Conversely, candidates with a Δ(*C*_*q*_) ≥ 3 cycles, indicating a degradation profile consistent with linear RNA species, were classified as unverified. This stringent validation criterion minimizes technical bias while providing robust confirmation of circRNA authenticity.

## Data availability

The circFormer model and the scripts for reproducing the analysis are available at [https://github.com/GenomicMedicine/circFormer].

## Supplemental data

Supplemental Tables 1-4; Supplemental Fig. 1.

## Declaration of generative AI and AI-assisted technologies in the writing process

During manuscript preparation, the authors used Claude and Perplexity to refine the language in some of their original writing and subsequently reviewed and edited the content as needed. The authors take full responsibility for the content of the work.

## Author contributions

WZ and SQ conceived, designed, and supervised the research. KL and WW designed and developed the circFormer approach and performed computational analysis. JJ, JD and JZ experimentally validated the results. KL, WZ, JJ and SQ analyzed the data. KL and WZ wrote the manuscript.

## Acknowledgments

The work was supported in part by funding from the Hong Kong RGC theme-based Strategic Target Grant Scheme (STG STG1/M-501/23-N), NSFC/RGC Collaborative Research Scheme (CRS_HKBU 2021/22), the Hong Kong RGC Collaborative Research Fund (CRF C5005-23WF), the Hong Kong Global STEM Professorship Scheme, and the Hong Kong Jockey Club Charities Trust.

---

**Supplemental Figure S1.**
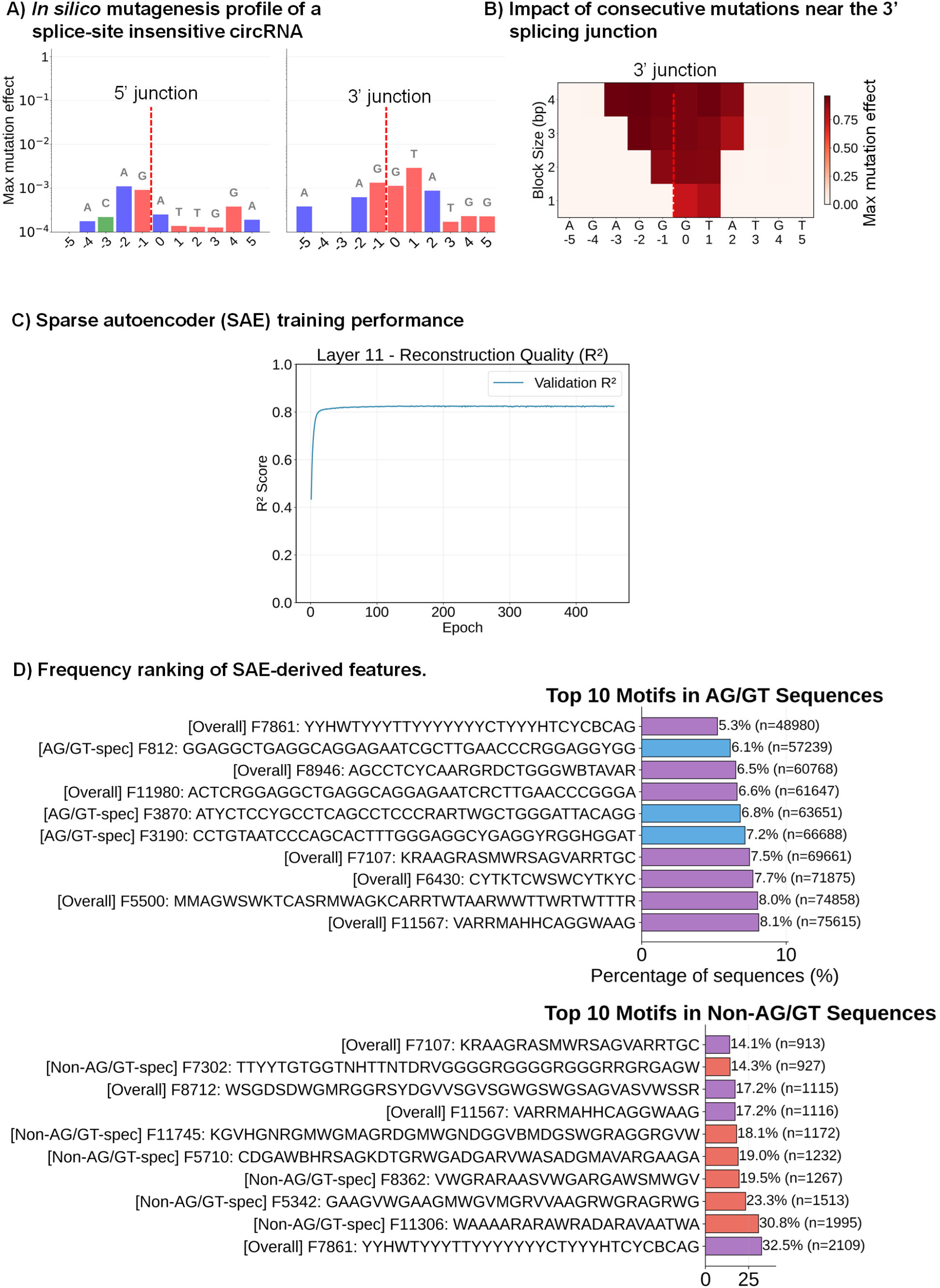
**A)** *In silico* mutagenesis profile of a splice-site-insensitive circRNA. Bar charts display the maximum mutation effects at the 5’ and 3’ back-splicing junctions for a representative true-positive AG/GT circRNA, showing minimal changes in prediction scores despite single-nucleotide mutations. **B)** Impact of consecutive mutations on model predictions. The heatmap shows the maximum mutation effect (color scale) resulting from mutating blocks of 1 to 4 nucleotides (y-axis) around the 3’ junction (x-axis), with darker red indicating greater impact. The effect of mutations near the 5’ back-splicing junction is not shown because their values are all less than 0.01. **C)** Reconstruction fidelity of the Sparse Autoencoder. The line plot shows the validation R-squared score (y-axis) as a function of training epochs (x-axis), assessing the SAE’s ability to reconstruct the original model embeddings. **D)** Prevalence ranking of discovered motifs in AG/GT and non-AG/GT circRNAs. Bar charts list the top 10 SAE-derived features ranked by frequency (x-axis) within AG/GT (top) and non-AG/GT (bottom) datasets. Purple, blue, and red bars represent, respectively, features shared overall, specific to AG/GT, and specific to non-AG/GT circRNAs.

